# Gpr19 is a circadian clock-controlled orphan GPCR with a role in modulating free-running period and light resetting capacity of the circadian clock

**DOI:** 10.1101/2021.08.07.455504

**Authors:** Yoshiaki Yamaguchi, Iori Murai, Kaoru Goto, Shotaro Doi, Huihua Zhou, Genzui Setsu, Hiroyuki Shimatani, Hitoshi Okamura, Takahito Miyake, Masao Doi

## Abstract

**Background and Purpose:** *Gpr19* encodes an evolutionarily conserved orphan G-protein-coupled receptor (GPCR) with no established physiological function in vivo. The purpose of this study was to determine the role of *Gpr19* in the circadian clock system.

**Experimental Approach:** We examined whether and how the master circadian clock neurons in the suprachiasmatic nucleus (SCN) express *Gpr19*. By analysing *Gpr19*-deficient (*Gpr19*^−/−^) mice, we asked whether *Gpr19* has a role in modulating free-running period and light resetting capacity of the circadian clock.

**Key Results:** Compared with the known common core clock genes, *Gpr19* was identified to show several distinct yet limited features related to the circadian clock. *Gpr19* mRNA was mainly expressed in the middle-to-dorsal region of the SCN. A conserved cAMP-responsive element within the *Gpr19* promoter drove the circadian expression of *Gpr19*. *Gpr19*^−/−^ mice exhibited a prolonged circadian period and a delayed initiation of daily locomotor activity in a 12-h light/12-h dark cycle. *Gpr19* deficiency caused the downregulation of several genes that normally peak during the night, including *Bmal1* and *Gpr176*. *Gpr19*^−/−^ mice had a reduced capacity for phase shift to early subjective night light. The defect was only observed for phase-delay, but not phase-advance, and accompanied by reduced response of c-Fos expression in the dorsal region of the SCN, while apparently normal in the ventral part of the SCN, in *Gpr19*^−/−^ mice.

**Conclusion and Implications:** *Gpr19* is an SCN-enriched orphan GPCR with a distinct role in circadian regulation and thus may be a potential target for alleviating circadian clock disorders.

- What is already known:

Gpr19 is an evolutionarily conserved class-A orphan receptor with no established physiological role in vivo.
The SCN is a light-entrainable master circadian pacemaker governing daily rhythms of behaviour and physiology.
- What this study adds:

*Gpr19* is an SCN-enriched orphan GPCR whose levels fluctuate in a circadian fashion.
*Gpr19* is a functional clock modulator involved in period determination and phase resetting.
- Clinical significance:

Targeting the orphan receptor Gpr19 may provide a therapeutic approach for alleviating circadian clock disorders.

## 1 INTRODUCTION

The SCN is the master circadian oscillator and the principal target for light modulation of the circadian rhythm in mammals (Herzog et al., 2017). Approximately 10,000 SCN neurons are clustered near the third ventricle above the optic chiasm, the source of direct retinal input to the SCN. The ventral part of the SCN close to the optic chiasm receives input from the retina, while the dorsal part of the SCN does not. Through communication between its ventral and dorsal parts, the whole SCN is synchronised to the ambient light/dark cycle (LeGates et al., 2014). The cyclic input serves solely to entrain the clock, not to sustain it.

The SCN generates endogenous circadian oscillation with a period (or time taken to complete a full cycle) of approximately 24 h. Animals, including human beings, can therefore sustain overt circadian oscillations in behaviour and physiology even under constant conditions, e.g. under constant darkness (Takahashi, 2017).

At the molecular level, individual neurons in the SCN act as cell-autonomous oscillators, exhibiting circadian oscillations of firing rate and gene expression. The rhythm-generating mechanism of the cellular clock involves clock genes, which regulate their own transcription in a negative transcription–translation feedback loop (TTFL). Positive regulators *Clock* and *Bmal1* and negative regulators *Per1*, *Per2*, *Cry1,* and *Cry2* constitute the main TTFL (Takahashi, 2017). Besides the clock components directly involved in the TTFL, SCN neurons express a number of genes that are involved in the coordination of cellular clocks within the structure. These are exemplified by VIP and its receptor Vipr2 coordinating the SCN circuit and expression of the circadian clock genes in the SCN (Aton et al., 2005; Colwell et al., 2003; Harmar et al., 2002). The AVP receptor V1a/b confers an intrinsic resistance against perturbation such as jet lag (Yamaguchi et al., 2013). The transcription factors Zfhx3 and Lhx1 regulate the expression of distinct neuropeptidergic genes to control circadian locomotor activity (Bedont et al., 2014; Hatori et al., 2014; Parsons et al., 2015). The G-protein signalling regulator RGS16 participates in circadian period determination by modulating cAMP signalling (Doi et al., 2011; Hayasaka et al., 2011). The orphan receptor Gpr176 also modulates the period of the SCN clock through circadian cAMP regulation (Doi et al., 2016). The neurotransmitter GABA has been implicated in synchronising individual cells within the SCN (Albus et al., 2005; Liu & Reppert, 2000). However, compared to the well-understood molecular mechanisms of the TTFL, molecular components involved in the coordination of the whole SCN are still not fully understood.

In the entrainment of the clock, phase resetting light pulses increase expression of *Per1* as well as other immediate early genes in the SCN. *Per1* induction changes the phase of the TTFL. In the SCN, indirect modulators of the TTFL also have a role in modifying the phase resetting response of the clock. Blocking the GABA_A_ receptor leads to increased phase shifts of circadian locomotor activity rhythm in mice (Lall & Biello, 2003). VIP-Vipr2 signalling is not only required for time keeping but is also involved in circadian clock entrainment to the environmental light-dark cycle (Hamnett et al., 2019; Hughes & Piggins, 2008; Mazuski et al., 2018; Patton et al., 2020). Lhx1 mutant mice rapidly phase shift under experimental jet lag conditions (Bedont et al., 2014; Hatori et al., 2014). Synaptic Ras GTPase-activating protein SynGAP and Ras-like G protein Dexras1 are involved in the modulation of light-induced phase shifts (Aten et al., 2021; Cheng et al., 2004). The voltage-gated channel Na_V_1.1 in the SCN is also required for the full phase-responsiveness of the clock (Han et al., 2012). These accumulating data support the notion of multilayered regulation of the capacity of phase response in the SCN clock, although the components involved are still not fully described.

*Gpr19* encodes an evolutionarily conserved orphan GPCR (https://www.gpcrdb.org/) first identified from a human genome EST library (O’Dowd et al., 1996). Histological studies previously identified the enrichment of *Gpr19* expression in the brain, including the SCN (Doi et al., 2016; Hoffmeister-Ullerich et al., 2004; Lein et al., 2007); however, how its expression is controlled in the SCN is not characterized. Moreover, currently, *Gpr19* lacks assignment to physiological functions; while a few published research articles reported on its potential association with certain metastatic cancers (Kastner et al., 2012; Rao & Herr, 2017; Riker et al., 2008), its distinct role in physiology has been unclear, reflecting the absence of study reporting the phenotype of *Gpr19*^−/−^ mice.

In the present study, we show that *Gpr19* is involved in the determination of the circadian period and phase-resetting capacity of the SCN clock. *Gpr19* mRNA was mainly expressed in the dorsal part of the SCN, with its expression fluctuating in circadian fashion. We explored the role for *Gpr19* in the regulation of circadian behaviour.

## 2 METHODS

### 2.1 Mouse strains and behavioural activity monitoring

*Gpr19*^−/−^ mice were obtained from the Mutant Mouse Resource & Research Centers (MMRRC strain name, *Gpr19^tm1Dgen^*) with a mixed genetic background involving 129P2/OlaHsd × C57BL/6J and backcrossed to C57BL/6J for ten generations prior to behavioural assessment. Single-caged adult male mice (8- to 15-week old) were housed individually in light–tight, ventilated closets within a temperature- and humidity-controlled facility. The animals were entrained on a 12-h light:12-h dark (LD) cycle at least 2 weeks and transferred to constant darkness (DD). Locomotor activity was detected with passive (pyroelectric) infrared sensors (FA-05 F5B; Omron) and data were analysed with ClockLab software (Actimetrics) developed on MatLab (Mathworks) (Doi et al., 2011). Free-running circadian period was determined with χ^2^ periodogram, based on animal behaviors in a 14-day interval taken 3 days after the start of DD condition. For phase shift experiments, mice were exposed to a 15-min light pulse at either CT6, CT14, or CT22 with a light intensity of 20 or 200 lux. Phase shifts were quantified as the time difference between regression lines of activity onsets before and after the light stimulation, using ClockLab software. All animal experiments were conducted in compliance with ethical regulations in Kyoto University and performed under protocols approved by the Animal Care and Experimentation Committee of Kyoto University (Approval No. 18-21-4). Animal studies are reported in compliance with the ARRIVE guidelines 2.0 (Percie du Sert et al., 2020) and with the recommendations made by the British Journal of Pharmacology (Lilley et al., 2020).

### 2.2 In situ hybridization

Radioisotopic *in situ* hybridization was performed as described with the following gene-specific probes (Shigeyoshi et al., 1997): for *Per1* (nucleotides 812–1651, NM_011065) and for *Gpr19* (nucleotides 923–1096, NM_008157). Free-floating brain sections (30-μm thick) containing the SCN were hybridized to anti-sense ^33^P-labeled cRNA probes. Quantification of expression strength was performed by densitometric analysis of autoradiograph films. To detect distribution of *Gpr19* mRNA expression in the SCN, RNAscope *in situ* hybridization was performed using 12 pairs of ZZ probe targeting the nucleotides 911–1583 of the mouse *Gpr19* (NM_008157). This region corresponds to the deleted sequence of the *Gpr19^tm1Dgen^* allele. The ZZ probes were designed and synthesized by Advanced Cell Diagnostics. RNA hybridization signals were visualized with the RNAscope 2-Plex Detection Kit (Advanced Cell Diagnostics) using the Fast Red chromogen according to the manufacturer’s protocol. Sections were counterstained with haematoxylin.

### 2.3 Immunoblot

Gpr19 antibody was raised in rabbit using a His-tag fused Gpr19 mouse protein fragment (amino acids 333–415). The raised antibodies were affinity-purified using a maltose-binding protein (MBP)-fused Gpr19 fragment (a.a. 333–415). Endogenous Gpr19 proteins were immunoprecipitated from the mouse hypothalamic SCN membrane extracts. The tissues were homogenized with a Dounce tissue grinder in a hypotonic buffer containing 20 mM HEPES (pH7.8), 2 mM EDTA, 1 mM DTT, and 1 × cOmplete Protease Inhibitor cocktail (Roche Diagnostics). After centrifugation at 20,400 × g for 30 min, the pellet was resuspended in a high-salt buffer containing 500 mM NaCl, 20 mM HEPES (pH7.8), 2 mM EDTA, 1 mM DTT, and 1 × cOmplete Protease Inhibitor cocktail. The mixture was then centrifuged, and the resultant pellet was solubilized with a detergent-containing buffer (20 mM HEPES [pH7.8], 150 mM NaCl, 2 mM EDTA, 1 mM DTT, 1% dodecyl-β-d-maltoside, 0.2% cholesteryl hemisuccinate, and 1 × cOmplete Protease Inhibitor). The soluble fractions, collected at either ZT4 or ZT16, were used for Gpr19 immunoprecipitation. Immunoblotting was performed using our standard method (Doi et al., 2011) with the same Gpr19 antibody.

### 2.4 5’ Rapid amplification of cDNA ends

Total RNA was purified from laser-microdissected mouse SCN using the RNeasy Micro Kit (Qiagen) according to the manufacturer’s instructions. The single strand cDNA for 5’RACE was prepared by in vitro reverse transcription with avian myeloblastosis virus reverse transcriptase XL (Takara Bio) using total RNA (0.5 µg) and the primer RT (5’-AGG ATG GAG GGA ATC-3’) and digestion of the template RNA with RNase H. 5’RACE was carried out using a 5’ Full RACE Core Set (Takara Bio). The first PCR was performed using the single strand cDNAs concatenated by T4 RNA ligase and primers S1 (5’-TTC TAT ACC ATC GTC TAC CCG CTG AGC TTC-3’) and A1 (5’-TTC AGC TCG TAC TGA AGC TCT GTC CTG TTG-3’) through a 25 cycle-amplification (94°C for 15 s, 45°C for 30 s, and 68°C for 2 min). Then, a nested PCR was applied to the first PCR products under the same condition using primers S2 (5’-GGG AAC TGC CTA TAC CGT CAT CCA CTT C-3’) and A2 (5’-CTC CTC ATG CAA TCC CAT CAG GCC ATG-3’). The resultant product of the nested PCR was cloned into pCR-Blunt II-TOPO vector for DNA sequencing.

### 2.5 Promoter activity reporter assay

We constructed a *piggyBac* (*PB*) transposon-based plasmid DNA containing *luciferase* (*luc*) reporter to ensure long-term transgene expression (Nakanishi et al., 2010). The following *PB*-based reporter plasmids were constructed: (i) pIR *Gpr19* promoter-*Luc2P* (−1083CREwt), in which a 1309-bp genomic DNA fragment of the murine *Gpr19* (−1083 to +226)-luciferase reporter (*luc2P*, Promega) was cloned into a vector engineered to contain the *PB* IRs and internal sequences necessary for efficient chromosomal integration (Nakanishi et al., 2010); (ii) pIR *Gpr19* promoter-*Luc2P* (−1083CREmut), which is the same as (i) except that the CRE was mutated to 5’-GCACAAAA-3’; (iii) pIR *Gpr19* promoter-*Luc2P* (−915CREwt), in which a 1141-bp genomic fragment of the *Gpr19* (−915 to +226) was cloned into the vector; (iv) pIR *Gpr19* promoter-*Luc2P* (−915CREmut), which is the same as (iii) except that the CRE was mutated to 5’-GCACAAAA-3’; (v) pIR *Gpr19* promoter-*Luc2P* (−514), which contains the −514 to +226 fragment of the *Gpr19*; and (vi) pIR *Gpr19* promoter-*Luc2P* (−242), which contains the −242 to +226 fragment of the *Gpr19*. For the analysis of isolated CRE activity, we used (vii) pGL4.23[luc2/minP] (Promega); (viii) pGL4.23 *Gpr19* 3×CREwt-*Luc2*, in which a tandem repeat of the sequence corresponding to the *Gpr19* CRE with its flanking sequences (positions −874 to −853) was cloned into the pGL4.23; and (ix) pGL4.23 *Gpr19* 3×CREmut-*Luc2*, which is the same as (viii) except that the CRE sequences were mutated to 5’-GCACAAAA-3’. All the plasmids were verified by DNA sequencing. MEF cells were uniformly plated in a 35-mm dish at a density of 3-4 × 10^5^ cells per dish and cultured for 1 day. Then, cells were transfected with a selected reporter plasmid using the Lipofectamine LTX/Plus reagent (Thermo Fisher Scientific). Where required, *PB* transposase-expressing vector (pFerH-PBTP) (Nakanishi et al., 2010) was co-transfected. Three days after transfection, culture medium was refreshed to the medium containing 1mM luciferin. On the following day, cells were treated with FSK (20 μM) or DMSO (1%). Luminescence was measured using a dish-type luminometer (Kronos Dio, ATTO). The average fold increase was determined by dividing the luciferase activity at 4−7 h post FSK or DMSO treatment with the average basal activity, which is 3-h reporter activity before FSK/DMSO treatment.

### 2.6 Viral transduction and bioluminescence recording of organotypic SCN slice culture

A luciferase reporter driven by a tandem repeat of the *Gpr19* CRE sequence (3×CRE-*Luc2P*) was inserted between the ITR sequences of pAAV-MCS vector (Cell Biolabs Inc) to obtain pAAV-3×CRE-*Luc2P*. HEK293T cells cultured in dish were co-transfected with pAAV-3×CRE-*Luc2P*, pAAV-DJ, and pHelper according to the manufacturer’s instructions (Cell Biolabs Inc). Three days after transfection, cells were harvested and resuspended in 1 ml of DMEM, followed by four freeze–thaw cycles and centrifugation. The titers of 3×CREwt-*Luc2P* and 3×CREmut-*Luc2P* virus solutions were ~8 × 10^12^ genome copies/mL. The SCN slices were prepared according to our standard method (Doi et al., 2019). Two days after the preparation of SCN slices, the AAV solution (3 μL per slice) was inoculated on the surface of the SCN slices. Infected slices were further cultured for ~14 days. Thereafter, luminescence from the culture was measured with a dish-type luminometer (Kronos Dio, ATTO) at 35°C using 1 mM luciferin (Doi et al., 2019). The luminescence was monitored for 2 min at 20-min intervals for each slice. The raw data were smoothed using a 1-h moving average and further detrended by subtracting a 24 h running average.

### 2.7 Laser microdissection and qRT-PCR analysis

Animals were sacrificed by cervical dislocation under a safety red light at the indicated time points in DD. Coronal brain section (30-μm thick) containing the SCN was prepared using a cryostat microtome (CM3050S, Leica) and mounted on POL-membrane slides (Leica). Sections were fixed in ice-cold ethanol-acetic acid mixture (19:1) for 2 min and stained with 0.05% toluidine blue. SCN were then excised using a LMD7000 device (Leica) and lysed into Trizol reagent (Invitrogen). Total RNA was purified using the RNeasy micro kit (Qiagen) and converted to cDNA with SuperScript VILO cDNA Synthesis kit (Invitrogen). qPCR was run on a BioMark HD System (Fluidigm) with a 48.48 Fluidigm BioMark Dynamic Array chip (Fluidigm) as described (Doi et al., 2019). The primer sets used for *Per1*, *Per2*, *Per3*, *Cry1*, *Cry2*, *Clock*, *Bmal1*, *Nr1d1*, *Dbp*, *E4bp4* and *Rplp0* were already reported elsewhere (Doi et al., 2019). The TaqMan probe and primers used for the other genes are listed in Table S1. The data were normalized with *Rplp0*. Hierarchical clustering was performed with Ward’s method by calculating Euclidean distances among the time-series data using scikit-learn (version 0.23.1) in Python. In this cluster analysis, the values of each mRNA expression were transformed by linear-scaling: the highest and lowest values were adjusted to 1 and 0, respectively. The resultant tree-diagram was further converted into an unrooted circular dendrogram, whose branch length reflects the degree of similarity between the genes, using the application Phylodendron software (http://iubio.bio.indiana.edu/treeapp/treeprint-form.html).

### 2.8 c-Fos immunolabeling

Free-floating immunohistochemistry was performed with 30-μm-thick serial coronal brain sections. To minimize technical variations in immunostaining, different tissue sections to be compared were immunolabelled simultaneously in a single staining mixture. c-Fos antibody (Abcam, ab7963, RRID:AB_306177, 1:10000 dilution) and biotinylated anti-rabbit IgG antibody (Vector Laboratories, BA-1000, RRID:AB_2313606, 1:1000 dilution) were used. Immunoreactivities were visualized with a peroxidase-based Vectorstain Elite ABC kit (Vector Laboratories) using diaminobenzidine as a chromogen. The number of c-Fos-positive cells in the SCN was counted with NIH ImageJ software. We used a rolling ball algorithm to correct uneven background in each photomicrograph. Nine SCN sections were examined per mouse. To measure c-Fos expression in the dorsal and ventral SCN, the SCN was divided into two regions in equal proportions along the vertical axis (from the dorsal-most to the ventral-most) for non-biased definition of the regions of interest. Three coronal SCN sections with characteristic dorsal and ventral subregions were used for counting.

### 2.9 Data and statistical analysis

The data and statistical analysis comply with the recommendations on experimental design and analysis in pharmacology (Curtis et al., 2018). All experiments were designed to generate groups of equal size, using randomization and blinded analysis. The statistical analysis was undertaken only for experiments where each group size was at least n = 5 of independent values and performed using these independent values. The group sizes for the in vivo experiments were chosen according to previous studies (Doi et al., 2016; Doi et al., 2019). Statistically significant differences between means of two groups were analysed by using unpaired Student’s *t*-test. For comparisons involving more than two groups, when F was significant, one-way or two-way ANOVA followed by Bonferroni’s *post hoc* test was performed. *P* values < 0.05 were considered significant. Outliers were included in data analysis and presentation. All statistical analysis was calculated using GraphPad Prism 8 (GraphPad Software, RRID:SCR_002798).

## 3 RESULTS

### 3.1 Expression of *Gpr19* in the SCN

We performed *in situ* hybridisation using a radioisotope-labelled probe for *Gpr19*. Coronal brain sections from wild-type (WT) mice confirmed the enrichment of *Gpr19* transcript in the SCN, while no signal was observed for *Gpr19*-deficient (*Gpr19*^−/−^) mice (Figure 1a). To detect distribution of *Gpr19* mRNA expression in the SCN, we next performed RNAscope *in situ* hybridization (Figure 1b). Coarse-grained RNA signals for *Gpr19* were mainly observed in the middle-to-dorsal region of the SCN in WT mice. Corresponding signals were not observed for *Gpr19*^−/−^ mice (Figure 1b). To test the possibility that *Gpr19* mRNA expression is regulated by the endogenous clock, we performed quantitative *in situ* hybridisation using samples from mice housed under constant dark conditions (DD). After entrainment on a regular 12-h light:12-h dark cycle (LD), mice were dark-adapted for 2 days before being sacrificed at 4-h intervals starting at circadian time (CT) 0 (Figure 1c, CT12 corresponds to locomotor activity onset). *Gpr19* mRNA was highest in the subjective day at CT4 and lowest in the subjective night at CT16, with an amplitude of ~2.75-fold (*P* < 0.05, CT4 vs CT16, one-way ANOVA with Bonferroni’s *post hoc* test, Figure 1c). These data demonstrate that the circadian clock regulates *Gpr19* expression in the SCN. We generated an anti-Gpr19 antibody. This antibody was unfortunately not useful for immunohistochemistry, but we confirmed its ability to specifically immunoprecipitate Gpr19 protein from WT mice but not *Gpr19*^−/−^ mice (Figure 1d). In this analysis we also noted a higher protein level of Gpr19 at daytime (ZT4, ZT represents Zeitgeber time; ZT0 denotes lights-on) than at night (ZT16) (Figure 1d). Thus, Gpr19 abundance appears to fluctuate at both mRNA and protein level.

**Figure 1.**
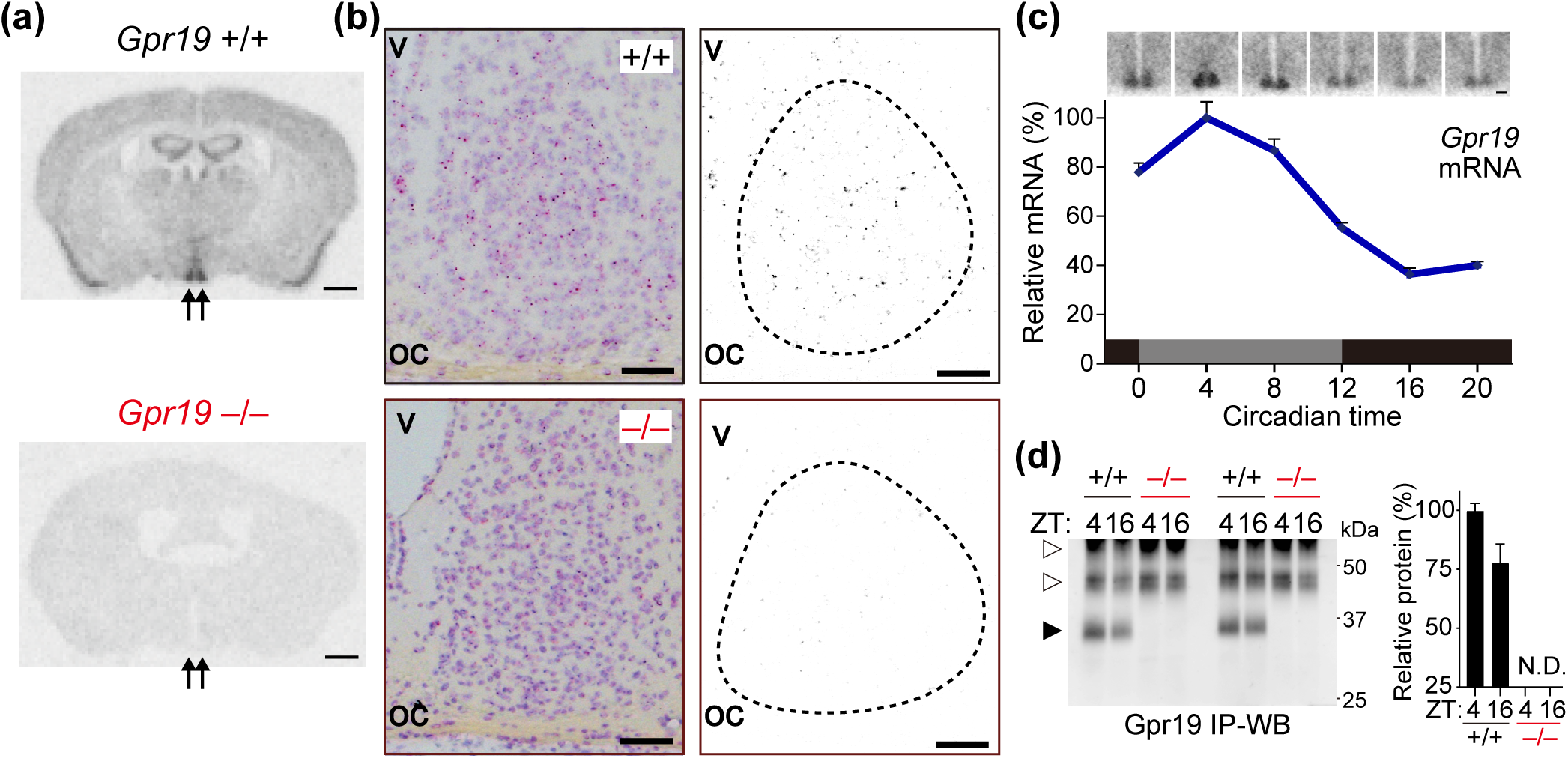
Spatiotemporal expression profile of *Gpr19* in the SCN. (**a**) Representative brain coronal sections of *Gpr19*^+/+^ and *Gpr19*^−/−^ mice hybridised to anti-sense ^33^P-labelled *Gpr19* riboprobe. Arrows indicate the position of the SCN. Scale bars, 1 mm. (**b**) RNAscope *in situ* hybridisation of *Gpr19* in the SCN. The sections were counterstained with haematoxylin. Right panels show the extracts of the *Gpr19*-RNAscope signal. The dashed lines delineate the SCN. oc, optic chiasm; v, third ventricle. Scale bars, 50 μm. (**c**) Circadian rhythm of *Gpr19* expression in the SCN. Relative mRNA abundance was determined by *in situ* hybridisation autoradiography. Values are presented as the mean ± SEM (*n* = 6, for each time point). Representative time-series autoradiographs are shown on top. Scale bars, 200 μm. (**d**) Western blots of Gpr19 in the SCN of *Gpr19*^+/+^ and *Gpr19*^−/−^ mice at ZT4 and ZT16. Endogenous Gpr19 proteins were immunoprecipitated from hypothalamic SCN membrane extracts and probed for Gpr19. The solid and open arrowheads indicate Gpr19 and non-specific bands, respectively. Relative protein levels were determined by densitometry.

### 3.2 CRE sequence in *Gpr19* promoter generates circadian oscillation in the SCN

To investigate the mechanism of circadian *Gpr19* expression, we performed sequence conservation analysis of the *Gpr19* promoter region among different mammalian species using the UCSC Genome Browser on Mouse (GRCm38/mm10, Figure 2). We identified the transcription start site by 5’ RACE using total RNA isolated from the SCN (Figure S1) and found a major site of initiation of *Gpr19*, which we designated as base pair +1 (Figure 2a). This analysis revealed two conserved segments, one of which was located near the transcription start site, including exon 1 (−194 to +232), while the other was located approximately 900-bp upstream of the gene (−1071 to −826). There were no consensus sequences matching the canonical circadian *cis*-elements E-box or D-box in these regions (Figure 2a). Instead, we identified a potential cAMP-responsive element (CRE) (−867 to −860) in the distal region. Of note, this conserved CRE sequence was functionally responsive, as revealed by the forskolin (cAMP enhancer)-dependent increase in reporter activity of *Gpr19* promoter-luciferase constructs that harbour the *Gpr19* CRE (−915CREwt and −1083CREwt, Figure 2b) but not of those with mutated CRE (−915CREmut or −1083CREmut) or shortened promoter constructs devoid of the CRE sequence (−514 and−242) (Figure 2b; see also Figure S2a−c). Vehicle treatment had no effect on the *Gpr19* promoter regardless of the presence of the CRE sequence (Figure S2d−f). Similar results were obtained with a reporter construct containing the isolated *Gpr19* CRE sequence (Figure 2a,b, 3×CREwt or 3×CREmut). With these results, we next moved to test whether the *Gpr19* CRE sequence is able to produce circadian transcriptional rhythm in the SCN. We performed long-term reporter recording using cultured SCN slices (Figure. 2c). Adeno-associated virus (AAV)-mediated 3×CREwt-luc expression in the SCN slice exhibited persistent circadian rhythms of bioluminescence over multiple cycles under constant culture conditions. In contrast, all tested slices expressing 3×CREmut-luc did not display detectable circadian luminescence expression (Figure 2c). The *Gpr19* CRE sequence, thus, has the ability to generate autonomous circadian expression in the SCN.

**Figure 2.**
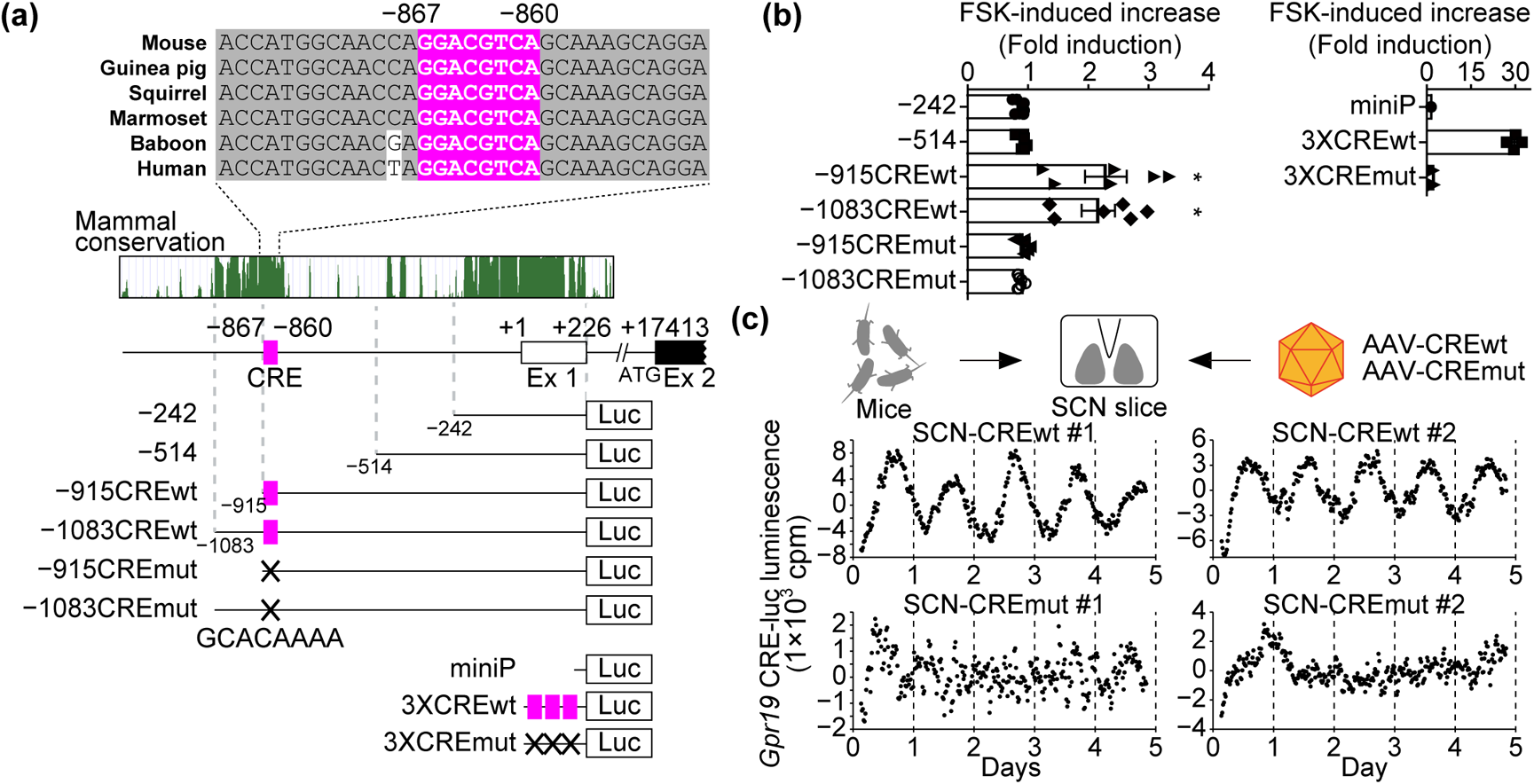
The *Gpr19* CRE sequence generates circadian expression of *Gpr19* in the SCN. (**a**) The CRE in the *Gpr19* promoter. Genomic positions relative to the transcription start site (+1) of the *Gpr19* gene are indicated along with evolutionary conservation scores among mammalian species. Alignment shows the CRE (−867 to −860; highlighted in magenta) and its flanking sequences of mouse, human, and other representative mammalian species. We used reporter constructs containing serial deletions of the mouse *Gpr19* promoter (−242 to +226, −514 to +226, −915 to +226, −1083 to +226) and the mutant derivative for the CRE (mut; GCACAA AA). We also used reporter constructs containing 3× isolated CRE (3×CREwt) or its mutant (3×CREmut). miniP, minimal promoter. (**b**) *Gpr19* promoter activities in MEF cells after treatment with cAMP enhancer FSK. Average fold increase relative to basal activity was calculated (*n* = 6, for each construct). Error bars indicate SEM. \**P* < 0.05, one-way ANOVA, Bonferroni’s *post hoc* test. (**c**) Representative detrended bioluminescence traces from SCN explants infected with AAV carrying the 3×CREwt (upper) or 3×CREmut (lower) reporter construct. Luminescence was recorded at 20-min intervals over 5 days in culture.

### 3.3 *Gpr19* deficiency lengthens the period of circadian locomotor activity rhythm

To assess the in vivo function of Gpr19, we monitored daily locomotor activity of *Gpr19*^−/−^ mice, which had been backcrossed to the C57BL/6J genetic background over 10 generations. Our initial survey using mice of a mixed background (75% C57BL/6J and 25% 129P2/ OlaHsd) suggested a trend towards prolonged periods of circadian locomotor activity for *Gpr19*^−/−^ mice compared to WT mice (free-running period (h), mean ± SEM; WT, 23.79 ± 0.04; *Gpr19*^−/−^, 23.92 ± 0.07, *P* = 0.1, *t*-test) (Doi et al., 2016). C57BL/6J-backcrossed mutant mice displayed an entrainment to a 12-h light:12-h dark (LD) cycle, although the phase of activity onset of *Gpr19*^−/−^ mice under LD conditions was delayed relative to that of WT mice (Figure 3a,b). On transfer of animals into constant darkness (DD), *Gpr19*^−/−^ mice showed a free-running period significantly longer than the WT period (WT, 23.77 ± 0.02; *Gpr19*^−/−^, 24.18 ± 0.03, *P* < 0.05, Student’s *t*-test, Figure 3c) and significantly longer than 24 h (95% confidence interval = 24.11, 24.25). These results indicate that Gpr19 is involved in the determination of circadian period length.

**Figure 3.**
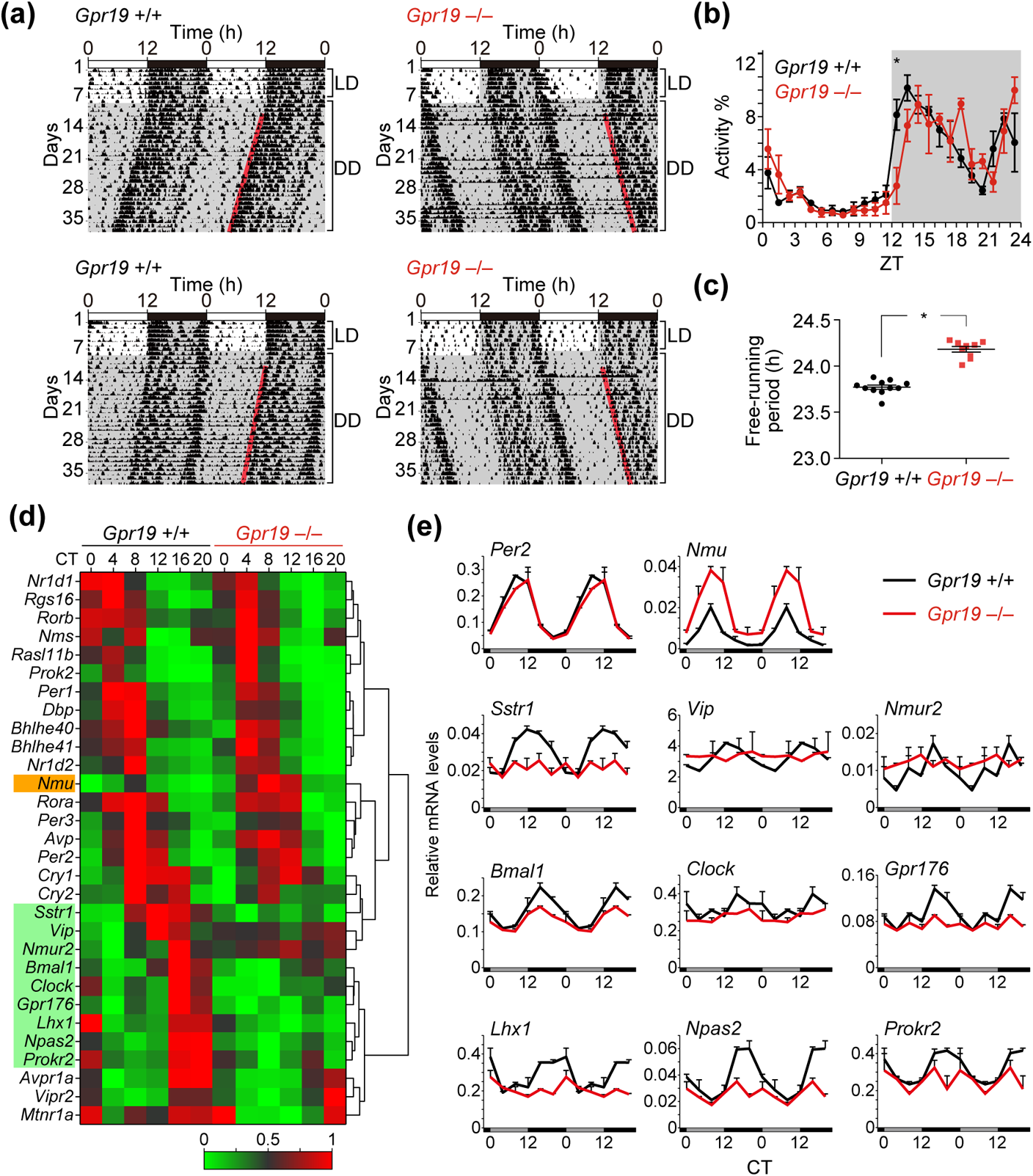
*Gpr19* deficiency elongates the period of locomotor activity rhythm and alters circadian clock gene expression in the SCN. (**a**) Representative double-plotted locomotor activity records of C57BL/6J-backcrossed *Gpr19*^+/+^ and *Gpr19*^−/−^ mice. Mice were housed in a 12L:12D light–dark cycle and transferred to DD. Periods of darkness are indicated by grey backgrounds. Each horizontal line represents 48 h; the second 24-h period is plotted to the right and below the first. The coloured lines delineate the phase of activity onset in DD. (**b**) Daily profile of locomotor activity of *Gpr19*^+/+^ and *Gpr19*^−/−^ mice in LD. Values are the mean ± SEM of % activity of a day. \**P* < 0.05, two-way ANOVA with Bonferroni’s *post hoc* test. (**c**) Period-length distribution of C57BL/6J-backcrossed *Gpr19*^+/+^ and *Gpr19*^−/−^ mice. Free-running period measurements were based on a 14-day interval taken after 3 days of a DD regime and were executed with a χ^2^ periodogram. Plotted are the period lengths of individual animals. Bars indicate the mean ± SEM (*Gpr19*^+/+^, *n* = 11; *Gpr19*^−/−^, *n* = 9). \**P* < 0.05, Student’s unpaired *t*-test. (**d**) Heatmaps displaying circadian expression of representative clock and clock-related genes in the SCN of *Gpr19*^+/+^ and *Gpr19*^−/−^ mice. The highest and lowest values of each gene were adjusted to 1 and 0, respectively. The genes (rows) are ordered by hierarchical clustering using Euclidean distance and Ward agglomeration. (**e**) Line graphs showing double-plotted circadian expression profiles of the genes affected by *Gpr19* deficiency in (**d**). Relative mRNA levels were determined by qRT-PCR (*n* = 2, for each data point). Values (mean ± variation) are double-plotted for better comparison between the genotypes. *Per2* is not affected. Data of all examined genes are shown in Figure S3.

### 3.4 *Gpr19* participates in maintaining proper circadian gene expression in the SCN

To identify potential molecular mediators of the effects of *Gpr19* deficiency in the SCN, we examined expression of representative clock and clock-related genes in the SCN of *Gpr19*^−/−^. The SCN of *Gpr19*^+/+^ and *Gpr19*^−/−^ mice housed in DD were micro-dissected every 4 h. Then, a customised panel of 41 SCN genes, which include representative core clock genes, clock-controlled genes, and circadian clock-related neurotransmitters and receptors, were analysed by quantitative RT-PCR using the Fluidigm system. The data of rhythmic genes were hierarchically aligned (Figure 3d) (see also plots in Figure S3). The core oscillatory gene *Per2* was basically circadian in the SCN of *Gpr19*^−/−^ mice, although the peak was slightly delayed, which was consistent with the prolonged circadian period of *Gpr19*^−/−^ mice (Figure 3e). Similarly, the genes with peak expression during daytime (e.g. *Per1*, *Cry1*, *Nr1d1*, *Rora*, *Bhlhe41*, *Prok2*, *Avp*, *Rasl11b*, and *Rgs16*) were apparently normal, except *Nmu*, whose expression was up-regulated in the SCN of *Gpr19*^−/−^ mice (Figure 3d,e; Figure S3). In contrast, a certain number of genes that show peak expression during the nighttime (CT12− 16) in WT mice, including *Bmal1*, *Clock*, *Npas2*, *Vip*, *Lhx1*, *Nmur2*, *Sstr1*, *Gpr176*, and *Prokr2*, were consistently downregulated in the *Gpr19*^−/−^ SCN (Figure 3d,e), suggesting that Gpr19 is involved in the maintenance of proper gene expression peaking in the night.

### 3.5 Gpr19 deficiency alters entrainment capacity

Next, we investigated the possible involvement of Gpr19 in entrainment of the clock. Light resets the phase of circadian rhythms in a phase-dependent and light-intensity-dependent manner, with delays dominating the early subjective night, advances dominating the late subjective night, and minimal phaseshifts during the subjective day. We illuminated mice with a short light pulse of 20 or 200 lux at CT14, CT22, or CT6 (Figure 4). In both WT and *Gpr19*^−/−^ mice, light at CT14 and CT22 caused the phase delay and advance, respectively, while light administered at CT6 had little effect (Figure 4). In addition, we observed that 200-lux light caused larger phaseshifts than 20 lux in both WT and *Gpr19*^−/−^ mice (Figure 4a,b). However, significant quantitative differences were detected in the magnitude of phase delays. Exposure to a 20-lux light at CT14 caused a delay of 1.90 ± 0.11 h in WT mice, whereas the phase-shifting response of *Gpr19*^−/−^ mice was only 0.71 ± 0.12 h (*P* < 0.05, two-way ANOVA with Bonferroni’s *post hoc* test) (Figure 4b,d). A 200-lux light pulse applied at CT14 resulted in a phase delay of 2.04 ± 0.12 h in WT and 1.16 ± 0.16 h in *Gpr19*^−/−^ mice (*P* < 0.05) (Figure 4a,c). In contrast, a light pulse given at CT22 led to comparable phase advances in WT and *Gpr19*^−/−^ mice at both 20 lux (0.36 ± 0.14 h for WT, 0.30 ± 0.09 h for *Gpr19*^−/−^) and 200 lux (0.72 ± 0.15 h for WT, 0.60 ± 0.16 h for *Gpr19*^−/−^). These results indicate that *Gpr19* is involved in determining the magnitude of phase delay of the circadian locomotor activity rhythm in mice.

**Figure 4.**
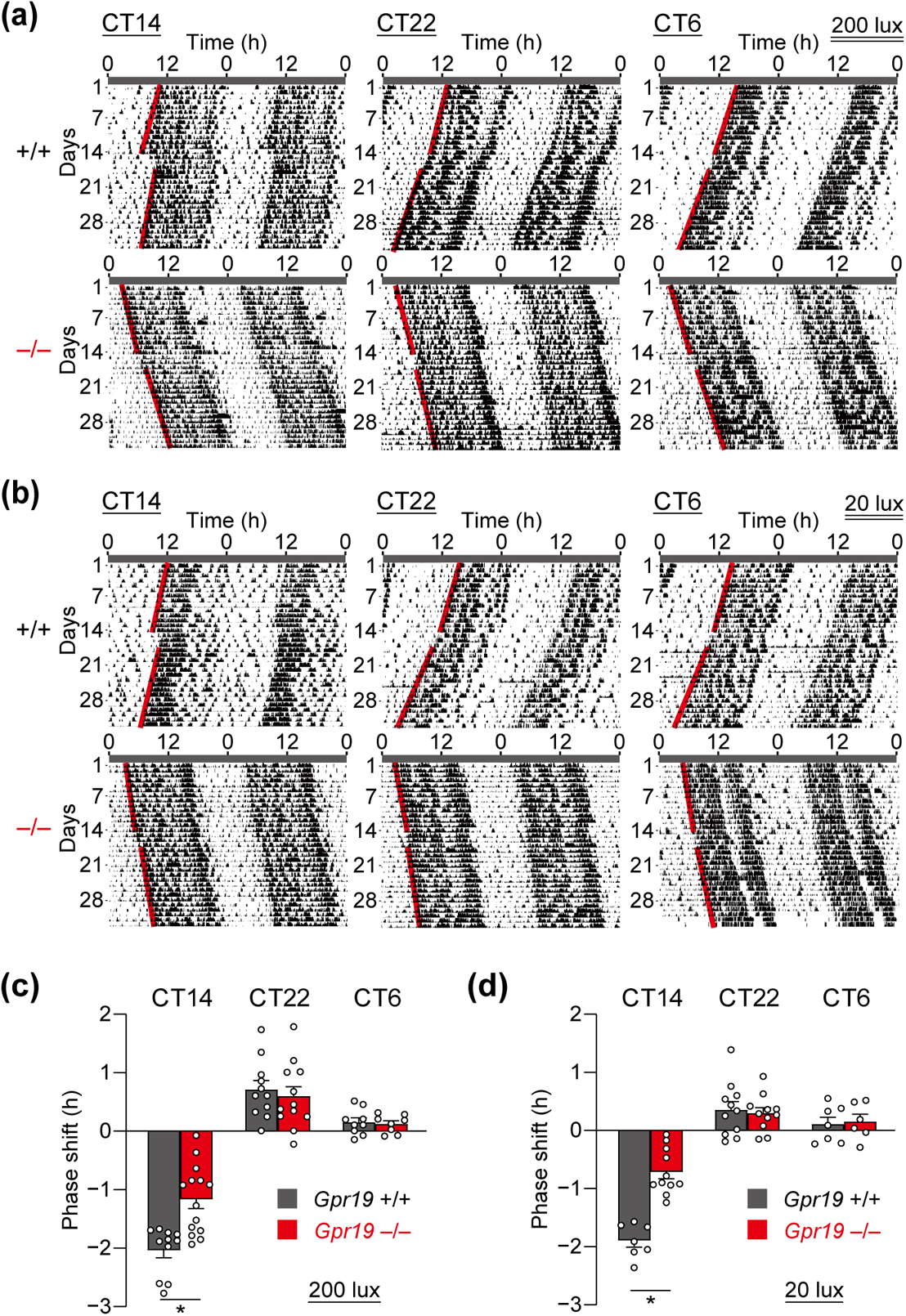
*Gpr19*^−/−^ mice exhibit a decreased capacity of phase shift to early subjective night light. (**a,b**) Representative double-plotted locomotor activity records of *Gpr19*^+/+^ and *Gpr19*^−/−^ mice before and after a 15-min light pulse exposure at CT14, CT22, or CT6. CT was determined for individual animals based on their free-running period and the onset of locomotor activity (which is defined as CT12). The red lines delineate the phase of activity onset. Phase shifts (delay at CT14, advance at CT22) were quantified as the time difference between regression lines of activity onset before and after the light pulse, 200 lux for (**a**) and 20 lux for (**b**). (**c,d**) Magnitude of light-induced phase-shifts of *Gpr19*^+/+^ and *Gpr19*^−/−^ mice. By convention, delays are negative, and advances are positive. Data indicate the mean ± SEM for 200 lux (**c**) and 20 lux (**d**) (200 lux: CT14, *Gpr19*^+/+^ *n* = 11, *Gpr19*^−/−^ *n* = 14; CT22, *Gpr19*^+/+^ *n* = 11, *Gpr19*^−/−^ *n* = 12; CT6, *Gpr19*^+/+^ *n* = 9, *Gpr19*^−/−^ *n* = 8; 20 lux: CT14, *Gpr19*^+/+^ *n* = 7, *Gpr19*^−/−^ *n* = 11; CT22, *Gpr19*^+/+^ *n* = 11, *Gpr19*^−/−^ *n* = 11; CT6, *Gpr19*^+/+^ *n* = 7, *Gpr19*^−/−^ *n* = 6). \**P* < 0.05, two-way ANOVA with Bonferroni’s *post hoc* test.

### 3.6 *Gpr19* deficiency alters light-evoked *Per1* and c-Fos expression in the SCN

To gain insight into decreased capacity of phase-shift of *Gpr19*^−/−^ mice, we examined the magnitude and location of *Per1* and c-Fos expression in the SCN of mice after light illumination. Mice were illuminated at CT14 or CT22 with 20 lux light, the intensity with which the difference in phase delay between WT and *Gpr19*^−/−^ mice was profound, and distribution of light-induced *Per1* and c-Fos expression in the SCN was examined either using radioisotopic *in situ* hybridization (for *Per1*) or immunohistochemistry (for c-Fos). *Per1* expression in the *Gpr19*^−/−^ SCN had a lower fold-induction ratio than that had in the WT SCN, at both CT14 and CT22 (CT14: 4.90 ± 0.09 for WT, 3.80 ± 0.04 for *Gpr19*^−/−^, *P* < 0.05; CT22: 3.60 ± 0.06 for WT, 3.15 ± 0.15 for *Gpr19*^−/−^, *P* < 0.05, two-way ANOVA with Bonferroni’s *post hoc* test), with apparently reduced *Per1*-positive-area in the SCN (Figure 5a). The number of c-Fos-immunopositive cells was decreased at CT14, but not CT22 (CT14: 999 ± 97 for WT, 625 ± 108 for *Gpr19*^−/−^, *P* < 0.05; CT22: 936 ± 224 for WT, 965 ± 160 for *Gpr19*^−/−^, *P* > 0.05, two-way ANOVA with Bonferroni’s *post hoc* test, Figure 5b). Within the ventral SCN, the increase in the number of c-Fos-positive cells was almost equivalent between the genotypes. Crucially, however, at CT14, the increased number of c-Fos-positive cells in the dorsal region was significantly reduced in the *Gpr19*^−/−^ SCN, compared to that in the WT SCN (c-Fos numbers in ventral: 286 ± 22 for WT, 245 ± 40 for *Gpr19*^−/−^, *P* > 0.05, in dorsal: 130 ± 15 for WT, 47 ± 7 for *Gpr19*^−/−^, *P* < 0.05, two-way ANOVA with Bonferroni’s *post hoc* test, Figure 5c−e), demonstrating an impaired expressional response of c-Fos in the dorsal part of the SCN in *Gpr19*^−/−^ mice.

**Figure 5.**
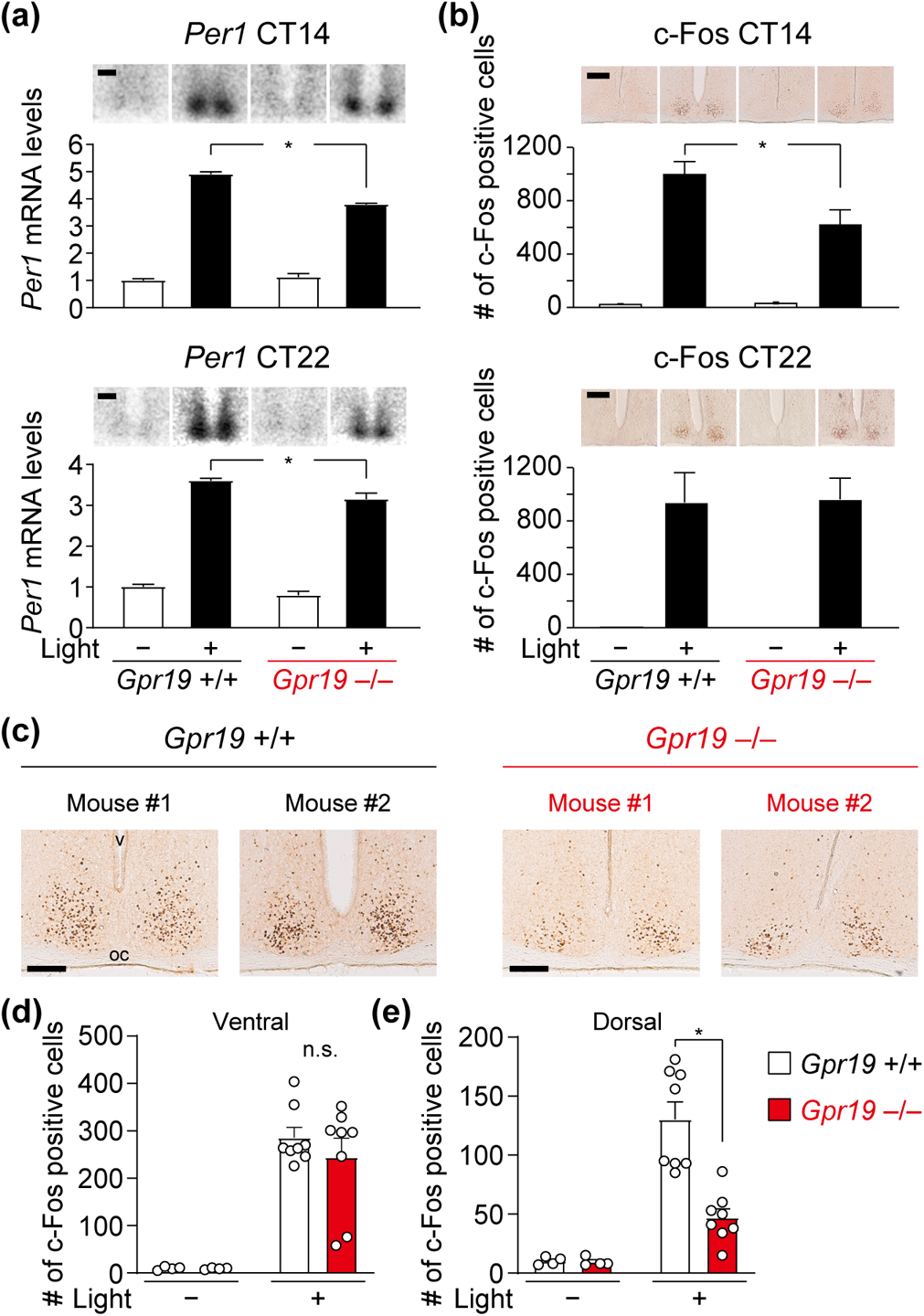
Attenuated light-induced induction of *Per1* mRNA and c-Fos immunoreactivity in the SCN of *Gpr19*^−/−^ mice. (**a**) *Per1* expression in the SCN of *Gpr19*^+/+^ and *Gpr19*^−/−^ mice with or without a 15-min light pulse exposure at CT14 or CT22. Mice were sacrificed 1 h after light onset. Data are presented as the mean ± SEM (*n* = 4). The mean value in *Gpr19*^+/+^ SCN without a light pulse was set to 1. \**P* < 0.05, two-way ANOVA with Bonferroni’s *post hoc* test. Representative autoradiographs are shown on the top. Scale bars, 200 μm. (**b**) The number of c-Fos-immunopositive cells in the SCN of *Gpr19*^+/+^ and *Gpr19*^−/−^ mice. Mice were illuminated as described in (**a**). Data are the mean ± SEM (*n* = 4 for light (−), *n* = 6−8 for light (+)). \**P* < 0.05, two-way ANOVA with Bonferroni’s *post hoc* test. Representative images of immunohistochemistry are shown on the top. Scale bars, 200 μm. (**c**) Reduced c-Fos induction in the dorsal area of the SCN in *Gpr19*^−/−^ mice. Representative images of immunohistological distribution of c-Fos expression in the SCN of *Gpr19*^+/+^ and *Gpr19*^−/−^ mice (2 mice for each genotype) after a 15-min light pulse exposure at CT14. oc, optic chiasm; v, third ventricle. Scale bars, 200 μm. (**d,e**) The number of c-Fos-immunopositive cells in the ventral (**d**) and dorsal (**e**) area of the SCN in (**c**) (*n* = 4 for light (−), *n =* 8 for light (+)). Values are the mean ± SEM. \**P* < 0.05, two-way ANOVA with Bonferroni’s *post hoc* test. n.s., not significant.

## 4 DISCUSSION

Besides clock components directly involved in the TTFL, SCN bears a number of additional genes implicated in modifying the length of circadian period and phase resetting capacity of the circadian clock (Herzog et al., 2017). A complete understanding of these additional modifiers of the SCN clock, however, still necessitate yet-unidentified related factors to be studied. In the present study, we demonstrate that the orphan G-protein coupled receptor *Gpr19*, whose mRNA expression exhibits circadian oscillation in the mid-to-dorsal region of the SCN, modulates the period and phase response of the circadian clock (a model, Figure 6).

**Figure 6.**
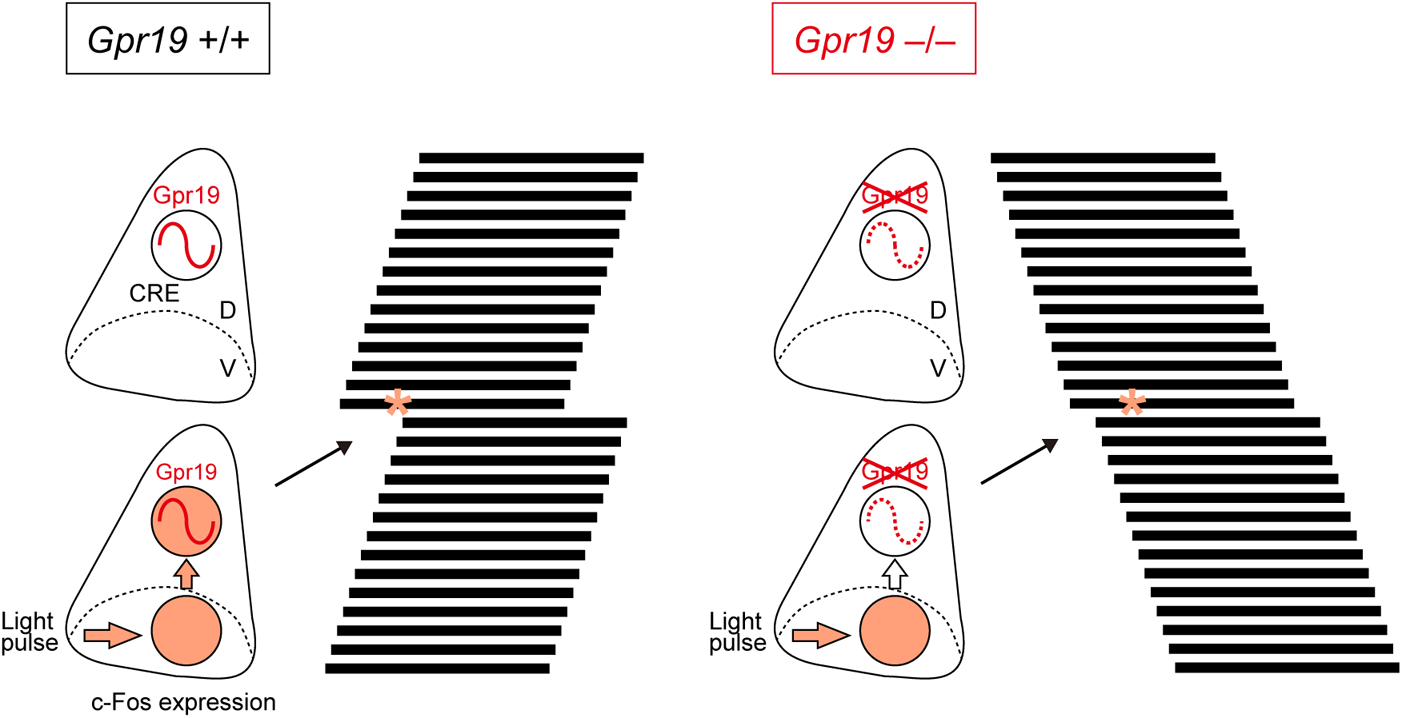
A putative role of Gpr19 in the central circadian clock modulation. The orphan receptor Gpr19 is a circadian oscillating GPCR localised to the middle-dorsal area of the SCN, is involved in the determination of the intrinsic period of locomotor activity rhythm, and modulates the extent of phase shift response to early subjective night light. Gpr19 controls gene expression in the SCN and modulates the propagation of light-entrainment signal from the ventral to the dorsal area of the SCN. Orange, c-Fos expression area; Asterisk, light pulse; D, dorsal area; V, ventral area; CRE, cAMP-responsive element.

We show that *Gpr19*^−/−^ mice exhibit a circadian period longer than 24 h under constant darkness. Under normal LD cycle conditions, these mice also show a delayed onset of locomotor activity compared to WT mice. The mechanism of this phase angle change is unknown, but a change from a circadian period shorter than 24 h to one longer than 24 h might be related to the observed phase angle phenotype of *Gpr19*^−/−^ mice (Johnson et al., 2003). A similar phase angle alteration was also reported in delayed sleep phase disorder patients (Micic et al., 2016) as well as several animal models, including Neuropeptide Y-deficient mice (Harrington et al., 2007), Na_V_1.1 channel mutant mice (Han et al., 2012), and lithium-treated mice (Iwahana et al., 2004).

Although the underlying molecular mechanism(s) of the lengthened circadian period of *Gpr19*^−/−^ mice is still unclear, we found a group of downregulated genes in *Gpr19*^−/−^ mice, the majority of which exhibit night-time peak mRNA expression in the SCN of WT mice. Thus, it is plausible to suggest that these alterations in gene expression may, at least in part, explain the phenotype of *Gpr19*^−/−^ mice. For example, *Bmal1* deficiency in SCN neurons has been previously reported to prolong the circadian period of locomotor activity rhythm (Mieda et al., 2015; Shan et al., 2020), consistent with the overall downregulation of *Bmal1* mRNA expression in the *Gpr19*^−/−^ SCN. *Clock*, *Npas2*, *Lhx1*, *Sst*, and *Gpr176*, which were also downregulated in the *Gpr19*^−/−^ SCN, are also involved in modulating the circadian period of locomotor activity rhythm (DeBruyne et al., 2007; Doi et al., 2016; Fukuhara et al., 1994; Hatori et al., 2014). The gene encoding Neuromedin U (Nmu) was, on the other hand, up-regulated in *Gpr19*^−/−^ mice, suggesting the possibility of a compensatory relationship between *Gpr19* and *Nmu*. These complex changes in mRNA expression of circadian clock-related genes might be part of mechanism explaining the phenotype of *Gpr19*^−/−^ mice.

A reduced magnitude of phase response to an early subjective night light pulse was also observed in *Gpr19*^−/−^ mice. In WT mice, a light pulse at CT14, of either 20 lux or 200 lux, caused a phase-delay of locomotor activity rhythm of approximately 2 hours. A reducing effect of the ablation of *Gpr19* on the magnitude of phase delay was more severe at a lower light-intensity condition: 20- and 200-lux light pulses caused phase delays of 0.71 and 1.16 h, respectively, in *Gpr19*^−/−^ mice. *Gpr19* is therefore likely to be required to induce the maximal phase delay response towards a light pulse of relatively low intensity.

Currently, we could not address the molecular mechanism of the reduced capacity of phase delaying in *Gpr19*^−/−^ mice. We observed that, in the *Gpr19*^−/−^ SCN, light-induced induction of *Per1* mRNA and c-Fos expression was attenuated in the dorsal region of the SCN. Thus, it is tempting to speculate that Gpr19 may function as an upstream regulator of *Per1* and c-Fos expression in the dorsal SCN. However, together with this interpretation, it can also be possible that Gpr19 may exert its indirect influence on the expression of *Per1* and c-Fos through affecting, for example, the gene expression required for the control of the circadian clock in the SCN. In this respect, the mRNA expression of *Lhx1* and *Sst*, both previously shown to play a role in circadian entrainment (Bedont et al., 2014; Hamada et al., 1993; Hatori et al., 2014), are downregulated in the SCN of *Gpr19*^−/−^ mice. It is also interesting to note that a similar ventral/dorsal phenotype, that is, a rather normal response in the ventral SCN but an impaired response in the dorsal SCN, has been previously described in Na_V_1.1 channel mutant mice (Han et al., 2012) and Sox2-deficient mice (Cheng et al., 2019). It is not known whether *Gpr19* has an association with these genes.

Our knockout study identified the role of orphan GPCR Gpr19 in the circadian clock system. In an attempt to identify its endogenous ligand, high-throughput ligand screening studies have been performed via several means, including Tango assay (Kroeze et al., 2015) and other β-arrestin recruitment-based assays (Colosimo et al., 2019; Foster et al., 2019). However, no cognate ligand has been determined for Gpr19 to date, hampering its further study in vivo using pharmacology. While adropin is considered a possible ligand for Gpr19 (Rao & Herr, 2017; Stein et al., 2016), its expression in the SCN has not been identified and the coupling between adropin and Gpr19 remains controversial (Foster et al., 2019). Apart from the SCN, *Gpr19* is also expressed in the testis, heart, liver, and kidney (Hoffmeister-Ullerich et al., 2004; O’Dowd et al., 1996) as well as certain cancer cell types (Kastner et al., 2012; Rao & Herr, 2017; Riker et al., 2008). The physiological role of Gpr19 in vivo, however, has not been well studied. Only a few published research articles suggest a role for Gpr19 in the regulation of cell cycle (Kastner et al., 2012) and MAPK signalling (Hossain et al., 2016; Thapa et al., 2018), using mRNA knockdown in in vitro cultured cells. In the present study, we provided the first report describing the role of Gpr19 in vivo, using *Gpr19* knockout mice. Our animal behavioural data demonstrate that Gpr19 is a functional component involved in the circadian clock. Pharmacological interventions targeting this orphan receptor may provide a potential therapeutic approach that modulates the circadian clock.

## ACKNOWLEDGMENTS

The authors thank Ichie Nishimura for technical support. This work was supported in part by research grants from the Project for Elucidating and Controlling Mechanisms of Ageing and Longevity of the Japan Agency for Medical Research and Development (JP20gm5010002), the Ministry of Education, Culture, Sports, Science and Technology of Japan (17H01524, 18H04015, 20B307), the Kobayashi Foundation, and the Kusunoki 125 of Kyoto University 125th Anniversary Fund.

## AUTHOR CONTRIBUTIONS

M.D. conceived the project; M.D. and H.O. designed the research; Y.Y., I.M., and K.G. performed experiments in collaboration with S.D., H.Z., G.S., H.S., and T.M.; M.D. and Y.Y. wrote the paper with input from all authors.

## DECLARATION OF TRANSPARENCY AND SCIENTIFIC RIGOUR

This Declaration acknowledges that this paper adheres to the principles for transparent reporting and scientific rigour of preclinical research as stated in the *BJP* guidelines for Design and Analysis, Immunoblotting and Immunochemistry, and Animal Experimentation, and as recommended by funding agencies, publishers, and other organizations engaged with supporting research.

## DATA AVAILABILITY STATEMENT

All data generated or analysed during this study are included in this published article (and its supplementary information files). The data that support the findings of this study are available from the corresponding author upon reasonable request.

