## Supplementary material for "Gpr19 is a circadian clock-controlled orphan GPCR with a role in modulating free-running period and light resetting capacity of the circadian clock": Figures S1-S5 Table S1

**Figure S1**

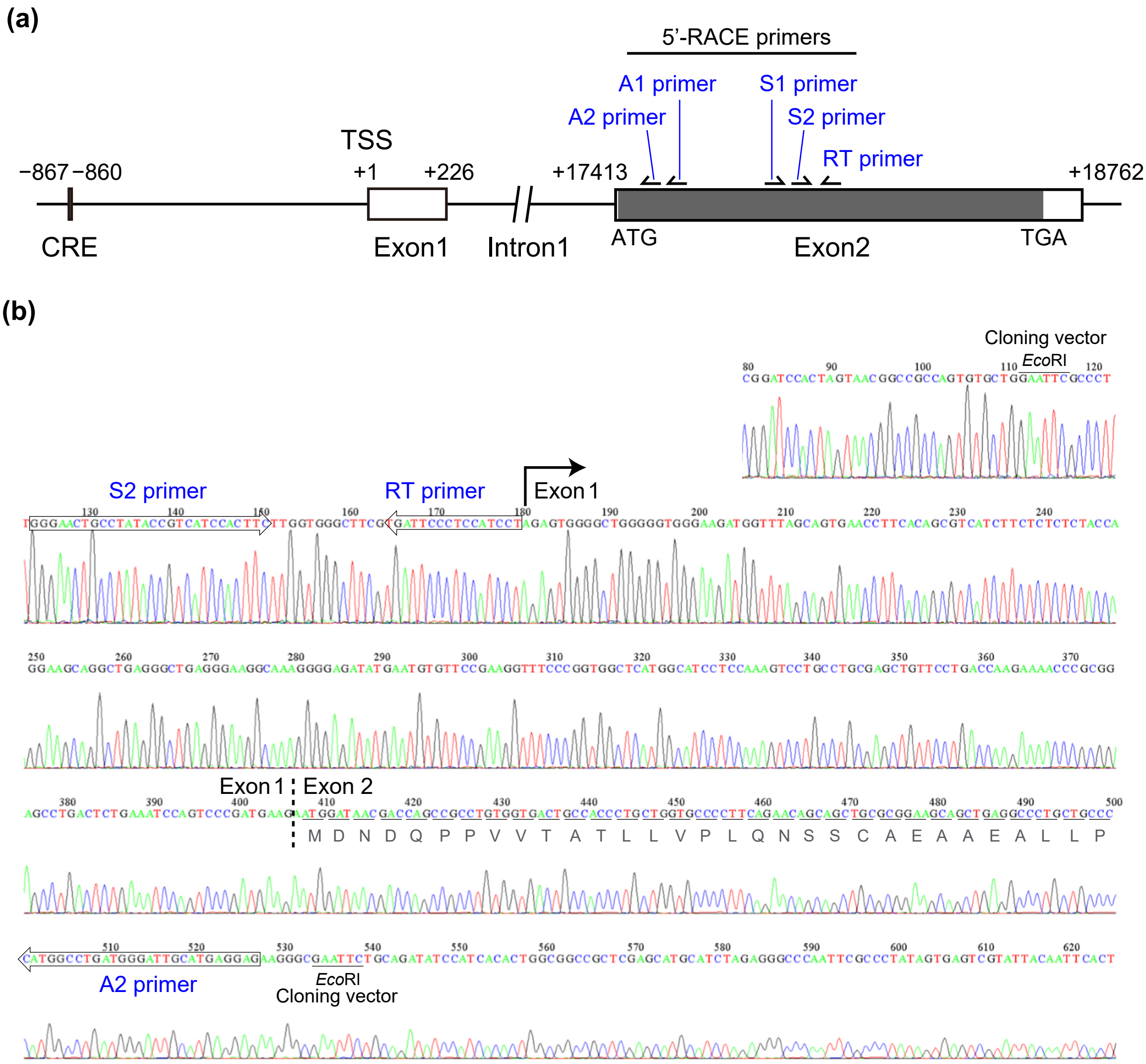


**Figure S1.** Determination of the transcription initiation site of *Gpr19* in the mouse SCN. (**a**) Schematic representation of the genomic organisation of the mouse *Gpr19*. Numbering shows the position relative to the transcription start site (TSS, +1) of *Gpr19* in the SCN. Open boxes represent exon 1 and exon 2. The exon 2 encodes the entire coding sequence of this gene. Arrows depict the 5′-RACE primers designed on the exon 2. (**b**) A representative DNA sequence chromatograph of the most common 5′-RACE fragment obtained from the mouse SCN. RNA extract from the mouse SCN was reverse transcribed with the RT primer. Single strand cDNAs were concatenated by RNA ligase and subjected to PCR with primers S1 and A1. Then, a nested PCR was applied to the first PCR products using primers S2 and A2. The resultant product of the nested PCR was cloned into pCR-Blunt II vector for DNA sequencing. Open arrows on the graph indicate the sequences of the 5′-RACE primers used. Bent arrow indicates the 5′-end of the exon 1. Vertical dashed line indicates the boundary between the exon 1 and exon 2. The exon 2 encodes expected Gpr19 amino acid sequence, which is indicated by single-letter code below the nucleotide sequence.

**Figure S2**

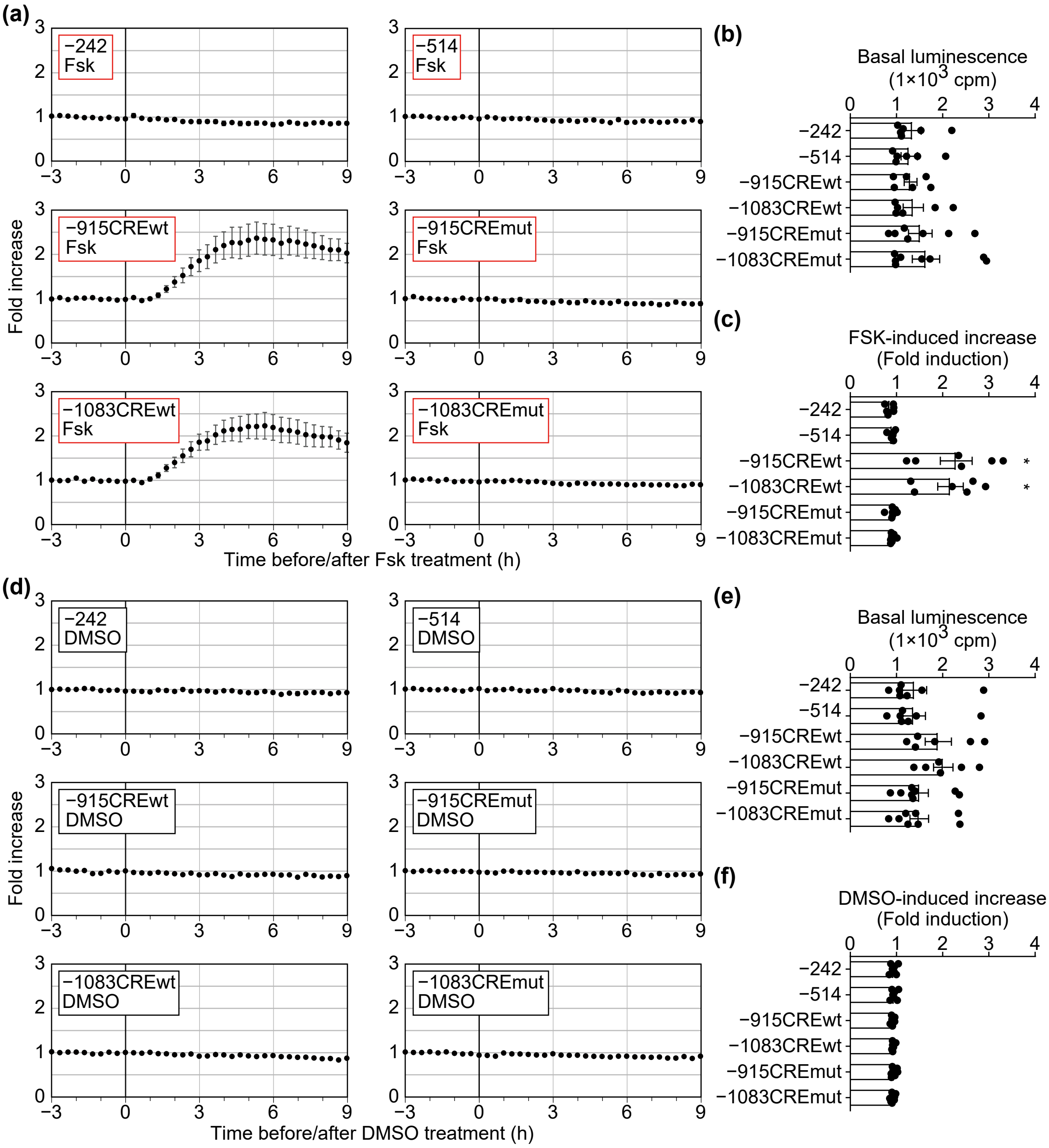


**Figure S2.** Reporter activity traces of WT and mutant *Gpr19* promoter fragments in MEF cells before and after FSK treatment, related to Figure 2b. (**a**) *Gpr19-luc2P* bioluminescence traces. Cells were treated with 20 µM FSK at time 0. Values are mean ± SEM ($n = 6$). (**b**) Average basal luminescence in (**c**). There was no statistically significant difference between the promoters in basal luciferase activity, determined using 3 h reporter activity before FSK treatment. (**c**) Average fold increase in luminescence by FSK in (**a**). Luciferase activity from 4 to 7 h post FSK treatment was divided by basal activity. The data are reproduced from Figure 2b. (**d**) *Gpr19-luc2P* bioluminescence traces in cells treated with vehicle (DMSO, 1%). Values are mean ± SEM ($n = 6$–$8$). (**e**) Basal luminescence in (**d**). (**f**) Fold change in luminescence after vehicle treatment in (**d**). Error bars in (**b**), (**c**), (**e**), and (**f**) indicate SEM. *$P < 0.05$, one-way ANOVA, Bonferroni's *post hoc* test.

### Figure S3

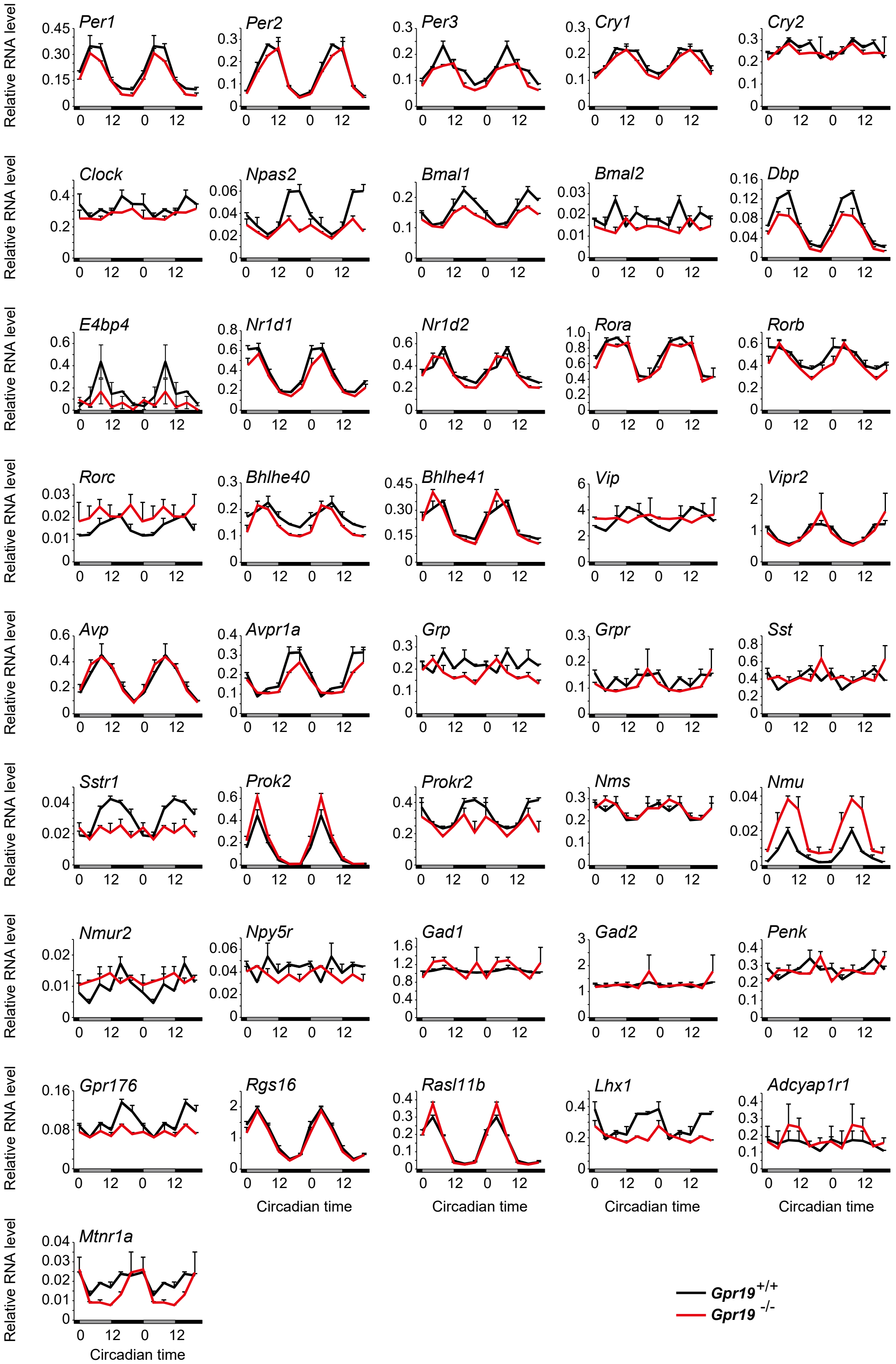


**Figure S3.** Circadian expression profiles of representative core clock genes, clock-controlled genes, circadian clock-related neurotransmitters and receptors in the SCN of *Gpr19*$^{+/+}$ and *Gpr19*$^{-/-}$ mice, related to Figure 3e. Relative mRNA levels were determined by qRT-PCR ($n = 2$, for each data point). Values (mean ± variation) are double-plotted for better comparison between the genotypes.

**Figure S4**

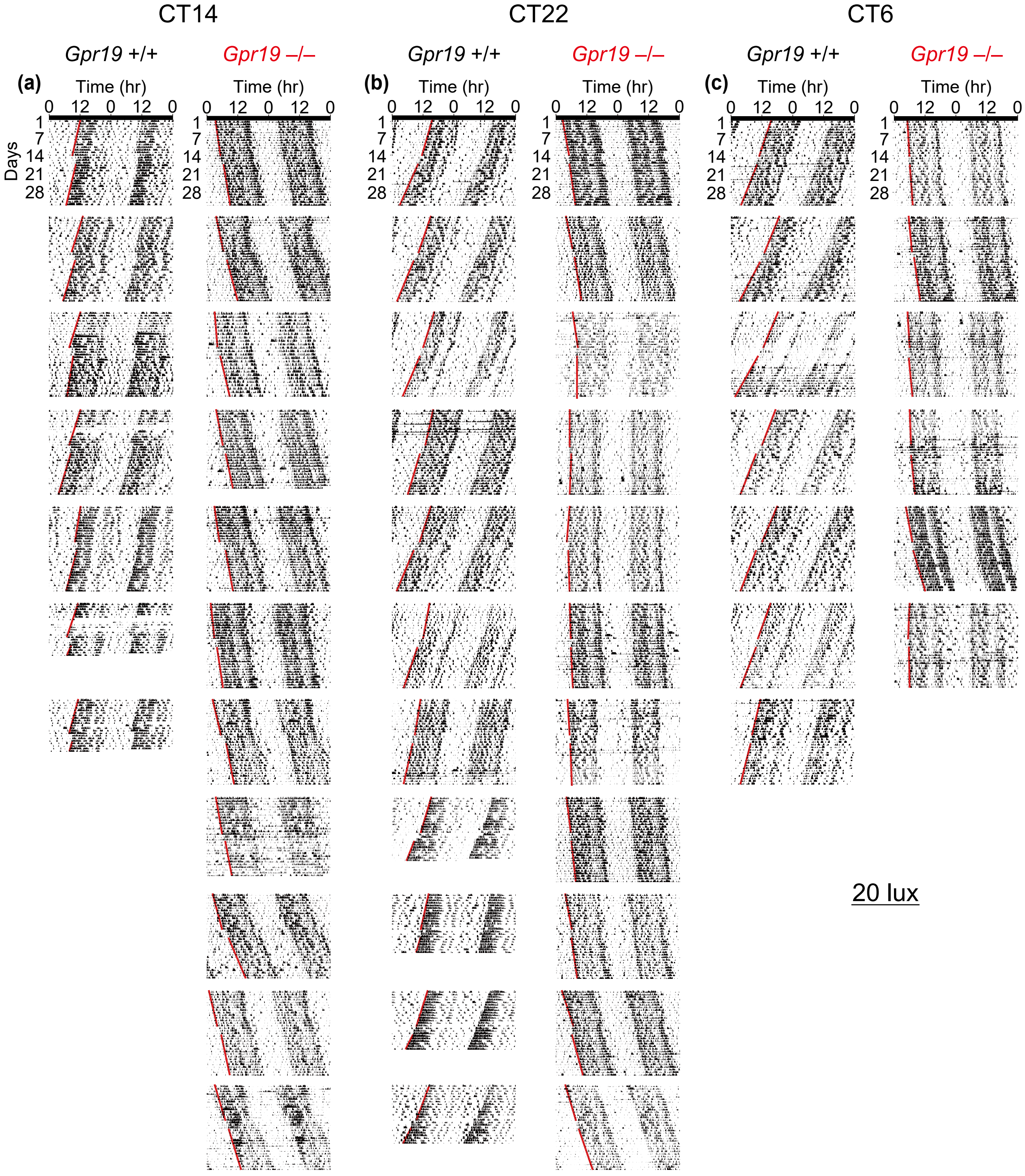


**Figure S4.** Circadian behavioral data of all individual mice before and after 20 lux light pulse exposure, related to Figure 4. Each panel shows double-plotted actograms of locomotor activity rhythms of *Gpr19*$^{+/+}$ and *Gpr19*$^{-/-}$ mice exposed 20 lux light pulse at (**a**) CT14, (**b**) CT22, or (**c**) CT6.

#### Figure S5

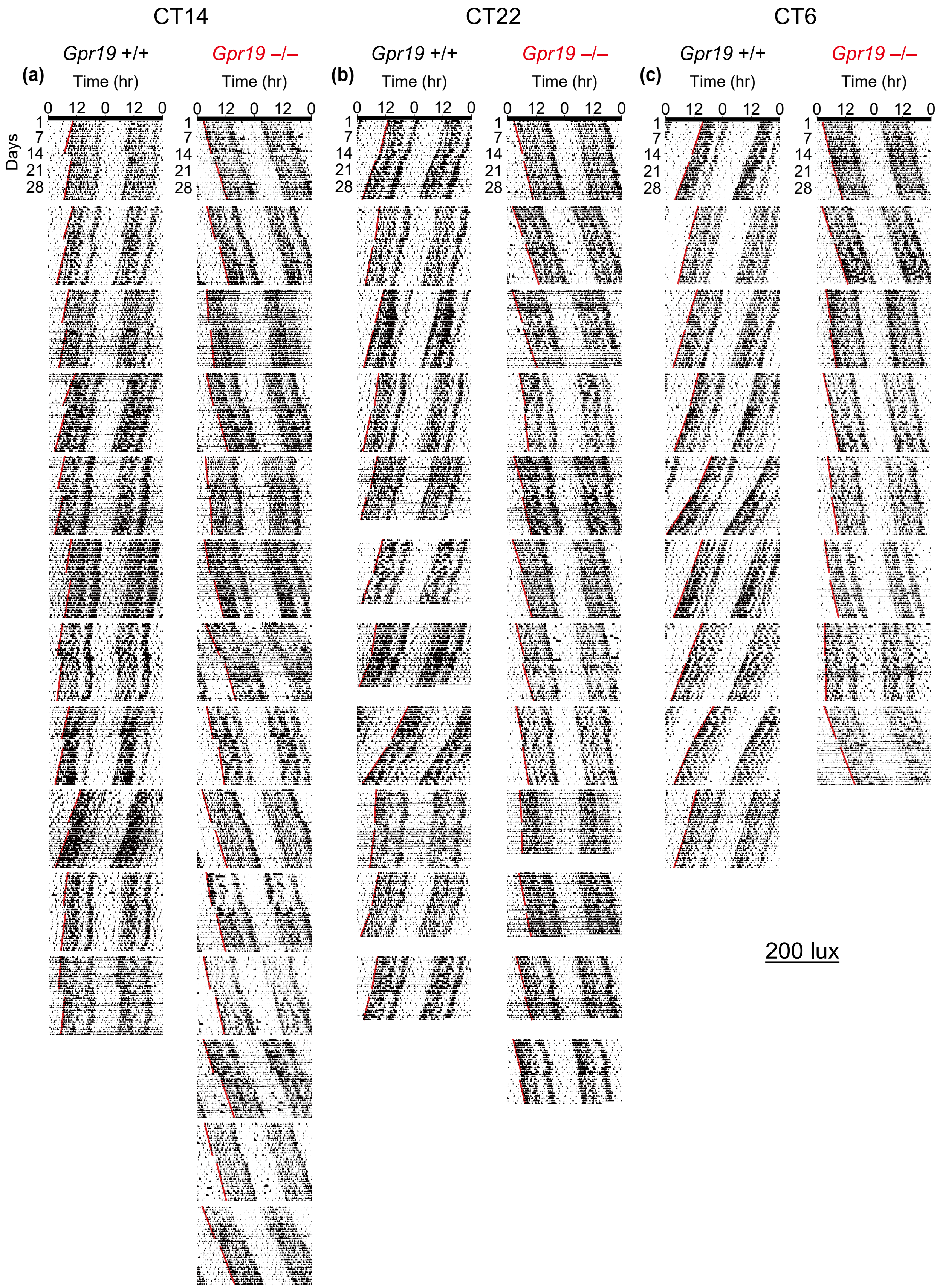


**Figure S5.** Circadian behavioral data of all individual mice before and after 200 lux light pulse exposure, related to Figure 4. Each panel shows double-plotted actograms of locomotor activity rhythms of *Gpr19*$^{+/+}$ and *Gpr19*$^{-/-}$ mice exposed 200 lux light pulse at (**a**) CT14, (**b**) CT22, or (**c**) CT6.

**Table S1.** TaqMan probes and primers

| Target gene | Probe (FAM-TAMRA) | Sense primer | Antisense primer |
|---|---|---|---|
| *Npas2* | TGGCTCTGTGCATACTTGACTTGCCGTC | CCAGCCCATCCAGCCTATGA | GCTGTTGGTAGGGTGTGAGTC |
| *Bmal2* | ACCACTTCCCGGTGACAGTGCCCA | AGCACTGAACCCGCCCAC | GCAGCCATGTCTATGCTGTCA |
| *Nr1d2* | CCACAGACAGACACTTCTTAAAGCGGCACT | TTCGGAGGAGCATTCAGCAA | CGAACAGCATCTCGTGACATC |
| *Rora* | TGGCAGAACTAGAACACCTTGCCCAGAA | GAGAGACTTCCCCAACCGTG | CTGGCAGGTTTCCAGGTGG |
| *Rorb* | TGATCGTTCTGACACAGCTCCATGAAGCCT | CGTGGTGGAGTTCGCCAAG | CTTCCAAGCAACCTGACTTCAGA |
| *Rorc* | CTGCGACTGGAGGACCTTCTACGGC | GTCTGCAAGTCCTTCCGAGAG | TCTCCCACATTGACTTCCTCTG |
| *Bhlhe40* | CCTCACGGGCACAAGTCTGGAAACCTG | AGCGGTTTACAAGCTGGTGAT | GGTCCCGAGTGTTCTCATGC |
| *Bhlhe41* | TCTTCTGATGCTGCTGCTCAGTTAAGGC | ACATCTGAAATTGACAACACTGGG | GCGCTCCCCATTCTGTAAAG |
| *Vip* | AGGGTCACCTGCTCCTTCAAACGGC | CCTTCTGTAGTGAGTAGGCTGGA | TCTGCAAGATGTCAGAGTCTGC |
| *Vipr2* | TGTCTCTGACCATCCATCGCTAGTGCA | CTACAGCAGACCAGGAAACAT | AGCCACACGCATCTATGAAATC |
| *Avp* | CGGCAAAGGACGCTGCTTCGGACC | CATGGAGCTGAGACAGTGTCTC | GGGCGAGGGCAGGTAGTTC |
| *Avpr1a* | CGATCACGGCGTTACTGGCTTCCTTGAAC | TGGACCGATTCCGAAAACCC | GCTCTGGACACAATCTTGTAGGA |
| *Grp* | AGCCACCAGCCACCTCAGCATCCG | AAGGGATTTGCTGGACCTCC | GAGTCTACCAACTTAGCGGTTTG |
| *Grpr* | TCCCACTGGCGATCATCTCTGTCTACT | ATGGCTTCCTTTCTGGTTTTCTAC | ATTCGATCTGCTTCTTGACATGTATA |
| *Sst* | CAAGGAAGATGCTGTCCTGCCGTCTCC | TAGACTGACCCACCGCGC | GGTGACACCGCCCAAAGC |
| *Sstr1* | TCATCAGTCCAGAGCCTTTCCACTTAATGG | CGATGGTGCCGCTTATTAATCA | TCCTGGGCACACTGGAGAG |
| *Prok2* | CCCGCCTTGCTGGTGGACCCA | CACCTTACTGTAGCATTGTGGG | GGGGATCTGGTTCAAACCTGTA |
| *Prokr2* | ACCAAGTAGGCAAGCCTCAACCAGAGC | GCGAGGAGAGCAGGACCAA | CAGGATTTGCCTCGGTGCTT |
| *Nms* | TCCACAATATCCAAGCCGTCGGGAGAATCA | CCCTCCTCAGGAGCTTCCC | CTGCTTCAGAAAGTATGCCAGTC |
| *Num* | TCTCCATCACTATACGGCAAAGCTCCCTCA | TGTGCGTCCTTTCTGTCCATTG | CTCACTTTGTTCCTGAGGCTTCTG |
| *Nmur2* | CTACATCCTCCCGATGACCCTCATCAGCG | CATCCAAGCTACCTCCTTCCTC | TCAGCCTGAGCCCCATGA |
| *Npy5r* | TCTCTGTGCCTCTGTAGTCCTCCCAGG | GAGGACTCTAGTATGGAGGTTAAAC | GTTCCGAGCAGCAGAAGTATTG |
| *Gad1* | ACCACCGAGCTGATGGCATCTTCCACT | ACCCTTGAACCGTAGAGACCC | ACACCAAGTATCATACGTTGTAGGG |
| *Gad2* | ACTGCTGCCATCCCCTTCTCCTTGACTT | GCTCATTGCCCGCTATAAGATG | GTGACTATGCTCTGATGTGAACG |
| *Penk* | AGGCAGCTGTCCTTCACATTCCAGTG | CGACATCAATTTCCTGGCGTG | GCAGGAGATCCTTGCAGGTC |
| *Gpr176* | ATGGTCAACTTGCCGCACAACAGTGTTCA | GCTCGCTACTGGGAAACTTCA | CACAGACCACACTGGCACAA |
| *Rgs16* | ATGAAGAAAGATTCCCAGCCGCGTCT | CCTGCCTGGAGAGAGCCAAA | CCCCAGTATCGGAGCTCAGC |
| *Rasl11b* | AGCCAGCGTTTCTCCTTCTATGTGAACTTG | CGAACGAAATGCAGGTAATCTCTAC | CGGAGTGTCCTGGACCTGAA |
| *Lhx1* | TTTGCAGCTACACCCAAGCCCACACG | CACGATCAAAGCCAAGCAACTG | AGACCTGGATAACACGCATGT |
| *Adcyap1r1* | TGCTGCCTATGGCTATTGCTATGCACTCTG | TCTCCCTGACTGCTCTCCTC | GCACATGGCTTGCTCCTTC |
| *Mtnr1a* | ACAGCCACCACGAGGTCTGCCACA | GAAGCTCAGGAACTCAGGGAATA | CCGTTGTTAAGGATAGATGTCAGC |
